# Transcranial magnetic stimulation of primary motor cortex elicits a site-specific immediate transcranial evoked potential

**DOI:** 10.1101/2023.09.05.556160

**Authors:** Mikkel Malling Beck, Lasse Christiansen, Mads Alexander Just Madsen, Armita Faghani Jadidi, Mikkel Christoffer Vinding, Axel Thielscher, Til Ole Bergmann, Hartwig Roman Siebner, Leo Tomasevic

## Abstract

**Background:** Transcranial evoked potentials (TEPs) measured via electroencephalography (EEG) are widely used to study the cortical responses to transcranial magnetic stimulation (TMS). Immediate transcranial evoked potentials (i-TEPs) have been obscured by pulse and muscular artifacts. Thus, the TEP peaks that are commonly reported have latencies that are too long to be caused by direct excitation of cortical neurons.

**Methods:** In 14 healthy individuals, we recorded i-TEPs evoked by a single biphasic TMS pulse targeting the primary motor hand area (M1_HAND_) or parietal or midline control sites. Sampling EEG at 50 kHz enabled us to reduce the duration of the TMS pulse artifact to a few milliseconds, while minor adjustments of the TMS coil tilt or position enabled us to avoid cranial muscular twitches during the experiment.

**Results:** We observed an early positive EEG deflection starting after approx. 2 ms combined with a series of peaks with an inter-peak interval of ∼1.1-1.4 ms in electrodes overlying the stimulated sensorimotor region. This multi-peak i-TEP response was only evoked by TMS of the M1_HAND_ region and was modified by changes in stimulation intensity and current direction.

**Discussion:** Single-pulse TMS of the M1_HAND_ evokes an immediate local multi-peak response at the cortical site of stimulation. Our results suggest that the observed i-TEP patterns are genuine cortical responses evoked by TMS caused by synchronized excitation of pyramidal neurons in the targeted precentral cortex. This notion needs to be corroborated in future studies, including further investigations into the potential contribution of instrumental or physiological artifacts.

## Introduction

Transcranial magnetic stimulation (TMS) can be combined with electroencephalography (EEG) to study cortical responsiveness to TMS [1]. A single TMS pulse targeting the motor hand representation in the precentral gyrus (M1_HAND_) elicits a series of positive and negative deflections in the averaged EEG waveform known as a transcranial evoked potential (TEP) [2]. The TEP peaks that are commonly reported occur at least 10 ms after stimulation [2]. Given their latencies, it is unlikely that these TEP peaks reflect the immediate cortical response evoked by inductive electrical stimulation in the M1_HAND_. Instead, conventional TEP peaks rather reflect a secondary excitation of cortical regions within the stimulated brain network, including the targeted M1_HAND_. The physiological interpretation of the standard TEP is further complicated by co-excitation of peripheral components of the nervous system causing peripherally evoked cortical responses [3,4]. TMS causes highly synchronized excitation of cortical pyramidal cells within the first few milliseconds after stimulation, potentially generating a measurable EEG signal [4]. Because of their very short latencies, immediate TEPs (i-TEPs) should not be affected by peripherally evoked cortical responses. So far, i-TEPs have not been in the focus of TMS-EEG research, because the cortical responses occurring at very short latencies have been obscured by co-activation of cranial muscles [5] and the TMS-pulse artifact [6–8]. The M1_HAND_ in humans and non-human primates has been shown to produce a characteristic series of descending volleys in the spinal cord in response to a single TMS pulse as revealed by epidural recordings [9–11]. These descending volleys are thought to be induced by the synchronized excitation of fast conducting pyramidal tract neurons in the M1_HAND_, and they leave the cortex already a few milliseconds after stimulation [11,12]. Given that this axonal activity in the descending pyramidal tract is evoked cortically, we hypothesized that TMS of M1_HAND_ may induce a synchronized patterned response in cortical pyramidal cells that can be captured in EEG electrodes overlying the stimulated M1_HAND_. We reasoned that the temporal pattern of pyramidal cell excitation would result in i-TEPs that mimic the temporal features of the corticospinal descending volleys. To test this hypothesis, we performed a TMS-EEG study that was optimized to detect short-latency i-TEPs in humans within a few milliseconds after single-pulse TMS. We therefore only tested participants in whom peri-threshold TMS of M1_HAND_ did not activate extracranial muscles after a tailored TEP-motor hot-spot hunting procedure. We further recorded EEG signals at high sampling rates (50 kHz) to reduce the duration of the stimulation artifact to approximately 2 ms.

## Methods

### Participants

Fourteen healthy volunteers (six males; age range: 23-32 years; mean age = 26.5±1.9 years) with no known neurological or psychiatric disorders participated in the experiments after giving written informed consent and being screened for MRI and TMS contraindications. The experiments were carried out in accordance with the Helsinki declaration and were approved by the regional ethical committee (Capital Region of Denmark; Protocol number H-15008824).

### Transcranial magnetic stimulation (TMS)

For all recordings, one-hundred biphasic TMS pulses were delivered using a figure-of-eight coil (MC-B70 coil, MagVenture X100 with MagOption, MagVenture A/S, Farum, Denmark) every 2 seconds (± 20% jitter) while participants were at rest. Biphasic waveforms were chosen as this allowed us to stimulate at lower absolute stimulation intensities due to lower cortical excitation thresholds [13], thus reducing the risk of evoking cranial muscle twitches [5]. During all experiments, the TMS coil positioning was monitored via stereotaxic neuronavigation (TMS Navigator, Localite, GmbH, Bonn, Germany) using individual T1w images (3T MRI, Siemens PRISMA). The recharge delay of the stimulator was set to 500 ms after the stimulation to avoid potential recharge artifacts in our signal of interest.

### Main experiment

The main experiment included 12 participants (5 males; age range: 23-32 years). Single biphasic TMS pulses were delivered over the left M1_HAND_ with intensities of 95% and 110% of individual resting motor threshold (RMT). The site of stimulation was first determined by finding the motor hand knob identified on individual structural MRIs [14]. Then, the coil was systematically moved in small steps until we found the site that elicited the largest and most consistent motor evoked potentials (MEPs). During this procedure, the coil handle was directed approximately 45 degrees lateral to the mid-sagittal plane to induce a current flowing perpendicular (anterior-posterior (AP) then posterior-anterior (PA)) across the precentral gyrus. We defined RMT as the stimulation intensity that elicited an MEP with a peak-to-peak amplitude of 50 µV in 5/10 consecutive stimulations expressed as percentage of maximal stimulator output (MSO). Next, we delivered ∼10 stimulations at 110% RMT to the motor hotspot while visually inspecting the averaged EEG data for scalp muscle artifacts [15] (in BrainVision Recorder Software). If this was the case, minor adjustments to the coil tilt or position were made in the medial direction, i.e., away from the m. temporalis and m. frontalis [5], and the above mentioned steps were repeated, i.e., RMT was re-estimated and EEG data was visually inspected. The coil position or tilt was modified in 6 of 14 subjects across all experimental conditions.

### Follow-up and control conditions

We further characterized the immediate EEG responses in a series of follow-up experiments to exclude the implication of known artifacts and verify that the observed i-TEPs may reflect genuine transcranial physiological responses.

### Site specificity

To investigate site-specificity of the i-TEP, we stimulated a posterior midline target site (N=9; 7 females) and a posterior lateral target (N=7; 5 females) in the posterior parietal cortex (PPC). The PPC site of stimulation was determined by standard Montreal Neurological Institute (MNI) coordinates (x = -26; y = -62; z = 56) obtained from NeuroQuery [16] projected into individual subject space. For the midline target, we stimulated between the CPz and the Pz electrode on the EEG cap and verified that this was at the midline using online neuronavigation. In both conditions, we ensured that the coil was tangential to the scalp and that the coil handle was pointing 45 degrees backward and lateral to the midsagittal plane. The stimulation intensity was set to 110% RMT_AP-PA_. In three subjects (2 females), we also adjusted the stimulation intensity for PCC stimulation to account for potential differences in scalp-to-cortex distance using Stokes’ formula [17]. In one male participant, we stimulated the left knee at 40% MSO corresponding to the average intensity used when stimulating at 110% RMT_AP-PA_ over M1_HAND_. In this experiment, 9 recording electrodes were attached in a 3×3 grid (approx. 1.5 cm apart) around the patella using double adhesive tape. The ground electrode was placed on the medial part of the tibial bone, while the reference electrode was placed on the head of the fibula.

## Effect of current reversal of TMS

In seven participants (5 females, age-range: 23-32 years), we stimulated M1_HAND_ with the current direction reversed (PA-AP) to assess how i-TEPs were modified by current direction. This was done at both 110% RMT_AP-PA_ and 110% RMT_PA-AP_ to include stimulation intensities at the same absolute and relative intensities, respectively.

### Intensity-response relationship

In four individuals (2 females), we obtained stimulus-response curves with both AP-PA and PA-AP current directions at stimulation intensities ranging from 60% to 160% of RMT_AP-PA_ in steps of 10% of RMT_AP-PA_. Two of the four subjects had participated in the main experiment. For measurements of stimulus-response curves, we used a smaller figure-of-eight coil with a 35mm diameter (Cool-B35, MagVenture A /S, Farum, Denmark) to further reduce the risk of eliciting cranial muscle twitches.

### EEG and EMG recordings

EEG was recorded in BrainVision Recorder software from 61 passive Ag/AgCl electrodes placed in an equidistant EEG cap (M10 cap layout, BrainCap TMS, Brain Products GmbH, Germany), with the reference and ground electrodes placed on the right and left side of the participants’ forehead, respectively. A TMS-compatible EEG amplifier (actiCHamp Plus 64 System, Brain Products GmbH, Germany) was used at a sampling frequency of 50 kHz (anti-aliasing low-pass filter at 10300 Hz) to minimize the duration of the TMS pulse artefact to ∼2 ms. All electrodes were prepared using electroconductive abrasive gel to reduce impedance (<5 kΩ), which was regularly checked during the experiment. Participants were equipped with modified earplugs playing pink noise with recorded TMS clicks generated using TAAC software [18]. The sound pressure was adjusted so that participants reported not being able to hear—or had largely attenuated—the click sounds generated by TMS while the coil was floating over the lateral surface of the head. Furthermore, a thin layer of foam (approx. 1.5 mm thick) was placed between the coil and the electrodes to reduce bone conduction of sound and vibration-induced somatosensory co-activation. During stimulation, participants were asked to keep their eyes open, avoid blinking, and fixate their gaze at a spot approximately 1m in front of them. Electromyographic (EMG) recordings were acquired from the right first dorsal interosseus (FDI) using a belly-tendon montage and a ground positioned on the right processus styloideus ulnae. EMG signals were amplified (x500), bandpass filtered (10-2000 Hz), and sampled at 2000 Hz (Digitimer D360, Digitimer Ltd., Hertfordshire, UK) using Signal software (v4.17, Cambridge Electronic Design, Cambridge, UK).

## Data analysis

EEG and EMG data were analyzed using custom MATLAB scripts (R2020a, v9.8, The MathWorks, Inc., Natick, MA, United States). Trials containing background EMG activity in a window 100 ms prior to stimulation were removed before averaging across epochs (median = 1; range: 0-35). These trials were also discarded from the EEG data. Peak-to-peak MEP amplitudes were individually computed and averaged within each condition to provide a measure of corticospinal excitability. Latencies for MEPs were also extracted and compared between stimulations with different current directions (Table S1). EEG data was epoched from -500 ms to 500 ms around stimulation and baseline corrected from -110 ms to - 10 ms. We observed a distinct high frequency (5000 Hz) ‘ringing’ response after the TMS pulse (lasting less than 2.5 ms) that was consistent across stimulation sites and current directions. This likely reflected residuals of the TMS pulse artifact and could be effectively removed with a band-pass filter (2nd order zero-phase Butterworth filter at 0.1-2000 Hz) (Figure S1). Filtered EEG data were visually expected for peak detection and extraction of the latencies and amplitudes of peaks and troughs in a window covering 2-8 ms after stimulation. To characterize the higher frequency peaks superimposed on the slower positive deflection, we calculated the difference between the first two peaks and the first trough. We used these relative amplitudes to illustrate the topographic distribution of the high-frequency peaks. Furthermore, we re-referenced to the average of all electrodes to investigate whether the findings were dependent on our recording montage (Figure S10).

## Results

### Immediate multi-peak EEG response to TMS of M1_HAND_

The 12 subjects who participated in the main experiment consistently showed an early EEG response to suprathreshold stimulation of the M1_HAND_ that occurred at very short latencies between 2 and 6 ms after the stimulus (Figure 1A). The i-TEP consisted of two to four positive peaks separated by short inter-peak intervals of 1.1−1.4 ms superimposed on a slower positive EEG deflection. The immediate multi-peak EEG response to TMS of M1_HAND_ was maximally expressed in electrodes located close to the pre-central site of stimulation (Figure 1A). Although i-TEPs were elicited in all subjects, there was some inter-subject variation in the number of observed peaks and their amplitudes and latencies (Figure 1B, 1C). The immediate cortical responses were followed by prototypical M1 TEP components with negative and positive deflections at well-known latencies, including the N15, P30, N45, P60, and N100 peaks (Figure S2).

**Figure 1.**
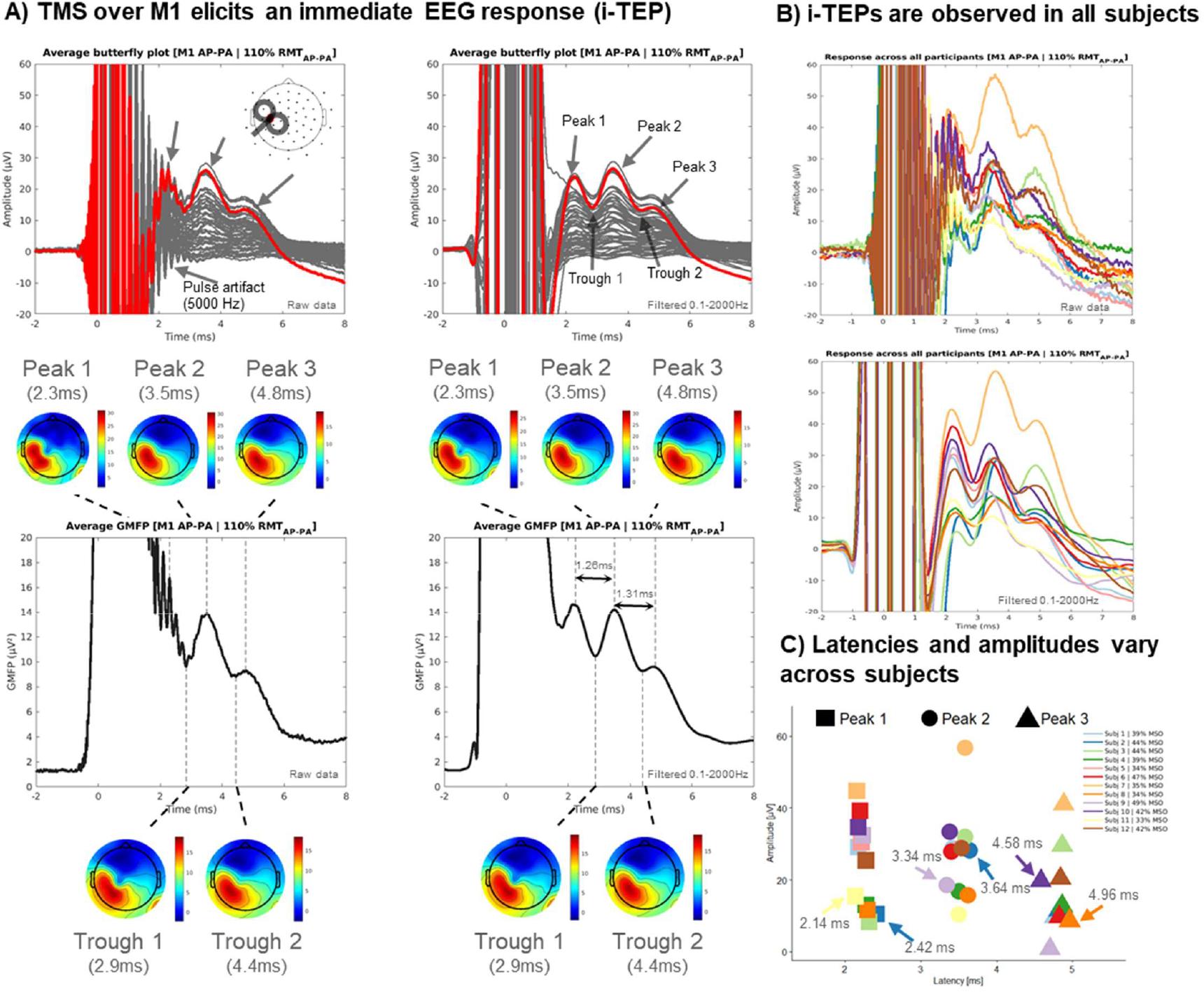
TMS over M1 elicits immediate TEPs (i-TEPs). Group-average EEG response to biphasic TMS (AP-PA) delivered at 110% RMT_AP-PA_ over M1_HAND_(N=12). Top row displays butterfly plots and bottom row the Global Mean Field Power (GMFP) with corresponding topoplots (min-max scaled). Left columns display raw EEG data whereas right column displays filtered EEG data (0.1-2000Hz, zero-phase, 2nd-order Butterworth filter; Figure S1). Trace highlighted in red corresponds to an electrode over left sensorimotor cortex close to the stimulating coil. TMS over M1 elicits a short-latency EEG response consisting of a series of peaks superimposed on top of a slower positive deflection. This response is largest in electrodes close to the site of stimulation in pericentral cortical regions. (B) displays the EEG response to TMS delivered at 110% RMT_AP-PA_ over M1_HAND_ from an electrode over sensorimotor cortex from all subjects in the main experiment (N=12). Following suprathreshold stimulation, i-TEPs are elicited in all subjects with some intraindividual variability in peak onset latencies and amplitudes. (C) Estimated individual i-TEP amplitudes and latencies following stimulation over M1_HAND_ at 110% RMT_AP-PA_ from band-pass filtered data. Minimum and maximum latency values per peak is highlighted to illustrate the variability of peak responses.

### Effect of stimulation intensity on i-TEP peak amplitudes

Even at a subthreshold stimulation intensity of 95% RMT_AP-PA_, we could still reliably evoke the first immediate peaks (Peak 1 and Peak 2) in all 12 participants. Compared to suprathreshold stimulation, immediate EEG responses evoked at subthreshold intensity were smaller in amplitude in all subjects, and Peak 3 was absent in 10 of 12 subjects (Figure 2; Figure S3).

**Figure 2.**
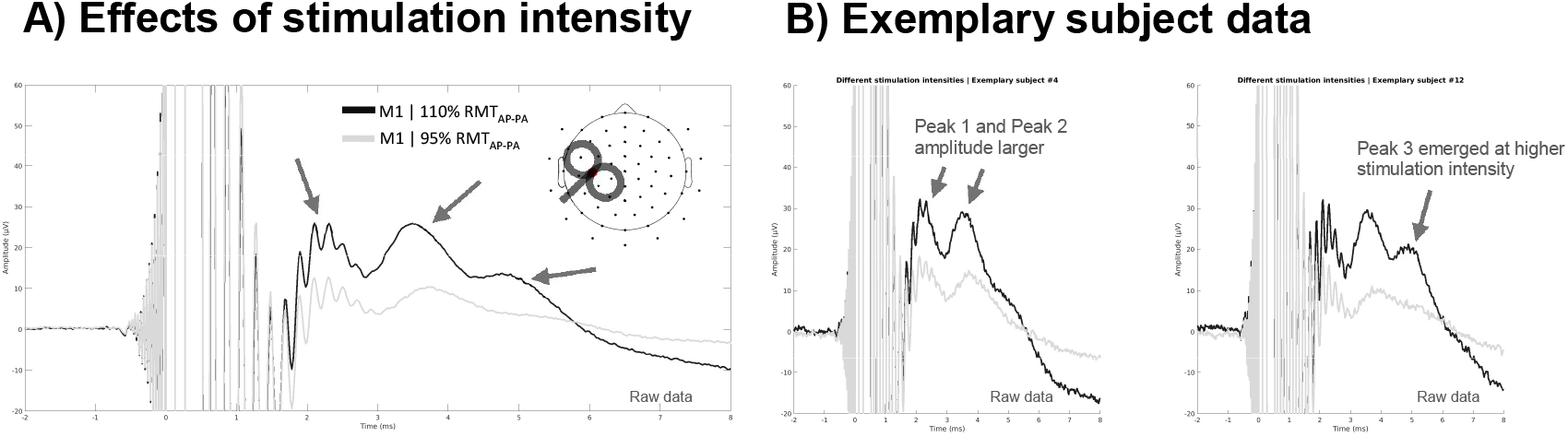
i-TEPs are modulated by stimulation intensity. (A) Group average EEG trace following biphasic TMS (AP-PA) delivered at 110% RMT_AP-PA_ (Black) and 95% RMT_AP-PA_ (light grey) over M1_HAND_ from an electrode over sensorimotor cortex (see electrode plot) from all subjects in the main experiment (N=12). (B) displays the EEG response to TMS at 95% RMT_AP-PA_ and 110% RMT_AP-PA_ in two exemplary subjects. Peak amplitudes of the i-TEPs scale with relative stimulation intensity (%RMT). This was observed in 12/12 subject (Figure S3). We also observed the appearance of a third peak following the 110% RMT_AP-PA_ condition (Peak 3). The third peak was clearly observed as a separate peak in 10/12 subject in the 110% RMT_AP-PA_ condition. The third peak was only clearly observed in 1/12 subjects in the 95% RMT_AP-PA_ condition (Figure S3).

### Site specificity of i-TEPs

The i-TEPs were only evoked when stimulating the M1 region (Figure 3, Figure S4 and Figure S5). TMS did not elicit i-TEPs or a comparable early response when stimulation was delivered to the PPC or a posterior midline target in any of the tested subjects (0/7 and 0/9 subjects, respectively). We adjusted the stimulation intensity to account for the larger scalp-to-cortex distance at the PPC stimulation site in three subjects, and while this affected the decay artefact, no i-TEP-*like* responses emerged (Figure S6). Stimulation of the left knee in one subject also did not yield any immediate responses in the recordings (Figure S7).

**Figure 3.**
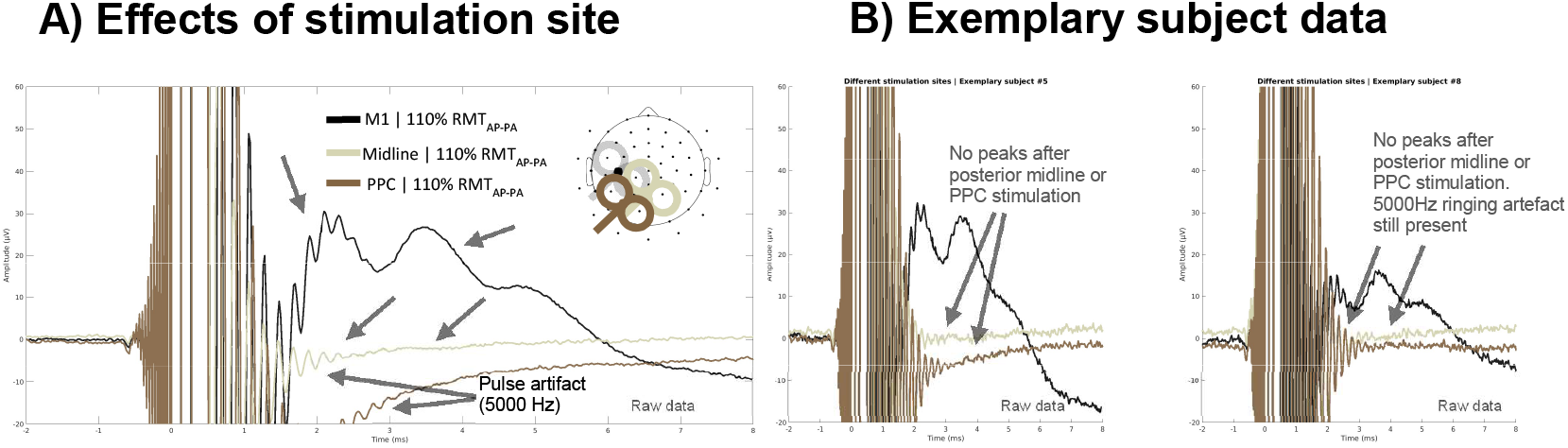
i-TEPs are specific to M1 stimulation. (A) group-averaged EEG response to biphasic TMS (AP-PA) delivered at 110% RMT_AP-PA_ to M1, as well as over one posterior medial (Midline; sand), and one posterior lateral control site (PPC, brown) in participants undergoing all three experimental conditions (N=7). Traces are extracted from electrodes close to the site of stimulation as shown in the electrode plot. (B) Individual unfiltered traces data from two exemplary subjects. i-TEPs are only observed after stimulating sensorimotor regions. We did not observe any immediate EEG responses after posterior midline (0/9) or PPC stimulation (0/7)(Figure S4; Figure S5).

### Effect of current reversal on i-TEP and MEP characteristics

Changing the direction of the TMS-induced current to PA-AP, as opposed to AP-PA, inverted the polarity of the TMS pulse artefact, but not the polarity of the i-TEPs (Figure 4A). However, current reversal did change the absolute and relative amplitudes of the i-TEP peaks. When using an identical stimulation intensity in terms of MSO, namely 110% RMT_AP-PA_, the observed evoked peak amplitudes for PA-AP were consistently smaller in all seven subjects (Figure 4BCD; Figure S8). Furthermore, Peak 1 was more difficult to discern and could only be clearly observed in 4 out of 7 participants (Figure 4BCD; Figure S8). In addition, when the stimulation intensity for the PA-AP was adjusted to individual RMT_PA-AP_, the amplitudes of Peak 2 and Peak 3 – but not Peak 1 – appeared larger in amplitude compared to AP-PA stimulation. In contrast, Peak 1 amplitudes were consistently greater for AP-PA stimulation (Figure 4DE; Figure S8). A similar behavior was observed for relative amplitudes where the troughs were subtracted from the peaks: For AP-PA stimulation, the early peak was greater and later peaks were smaller with respect to PA-AP current direction and vice versa. Taken together, these observations suggest that changing the direction of the induced current to evoke anterior-to-posterior (AP) currents facilitated the later i-TEP peaks.

**Figure 4.**
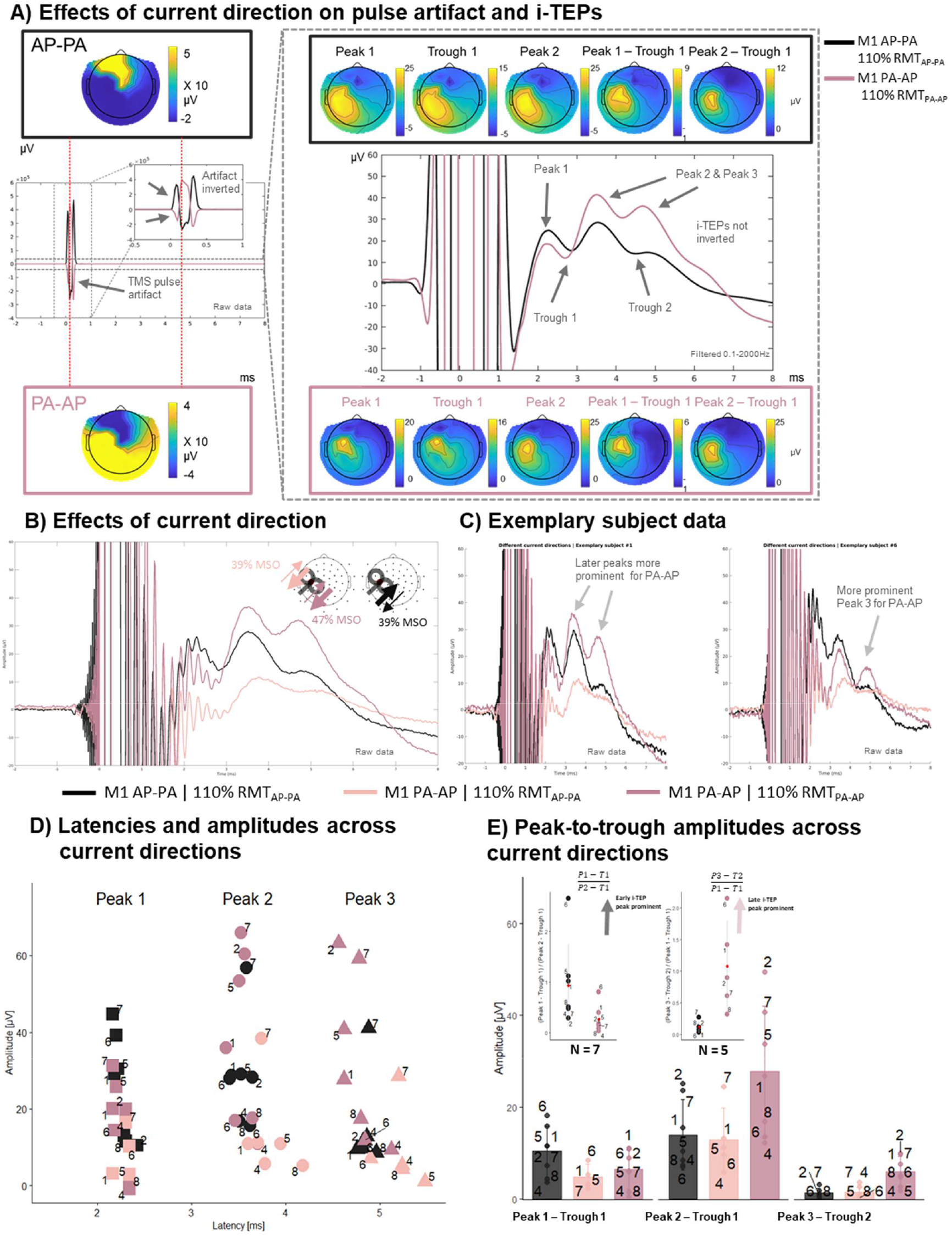
i-TEPs are influenced by current direction. (A) Left displays the raw group-averaged EEG response to biphasic TMS with AP-PA current direction delivered at 110% RMT_AP-PA,_ and PA-AP stimulation at 110% RMT_AP-PA_ and 110% RMT_PA-AP_ over M1_HAND_ (N=7). Traces are extracted from an electrode close to the site of stimulation. Note that the y-axis is scaled to capture the entire TMS pulse artifact. Right part of the figure displays group-averaged filtered EEG data responses and their topography for the same stimulation conditions. Note that the y-axis is scaled to capture the i-TEP responses. The polarity of the pulse artifact is inverted when reversing the stimulation current whereas this is not the case for the i-TEPs. (B) displays group averaged i-TEP responses for AP-PA at 110% RMT_AP-PA_ and PA-AP stimulation at 110% RMT_AP-PA_ and 110% RMT_PA-AP._ (N=7)(C) shows the same conditions for two exemplary subjects (for remaining subjects see Figure S8). i-TEPs display different quantitative and qualitative features based on used current direction. (D) displays the individual peak amplitudes and latencies for the different stimulation conditions for each subject (label reflects subject number) and (E) displays differences in peak-to-trough amplitudes across conditions. The inset displays the relative weighting of early vs. late i-TEP peaks. AP-PA stimulation leads to greater early i-TEP amplitudes, whereas PA-AP stimulation results in greater late i-TEP amplitudes.

### Dose-response relationship

We explored changes of i-TEPs and MEPs when changing stimulation intensity by characterizing intensity-response relationships in four subjects for both AP-PA and PA-AP stimulation (Figure 5; Figure S9). Both i-TEPs and MEPs gradually increased with the intensity of TMS. In two subjects, we observed that i-TEPs emerged around active motor threshold (AMT), and in two individuals i-TEPs were observed between AMT and RMT of the respective current directions. Using PA-AP currents, i-TEPs and MEPs were elicited at higher absolute stimulation intensities in 3 of 4 individuals (Figure 5; Figure S9). Together, the results showed that i-TEPs appeared at lower absolute stimulation intensities following AP-PA stimulation compared to PA-AP stimulation.

**Figure 5.**
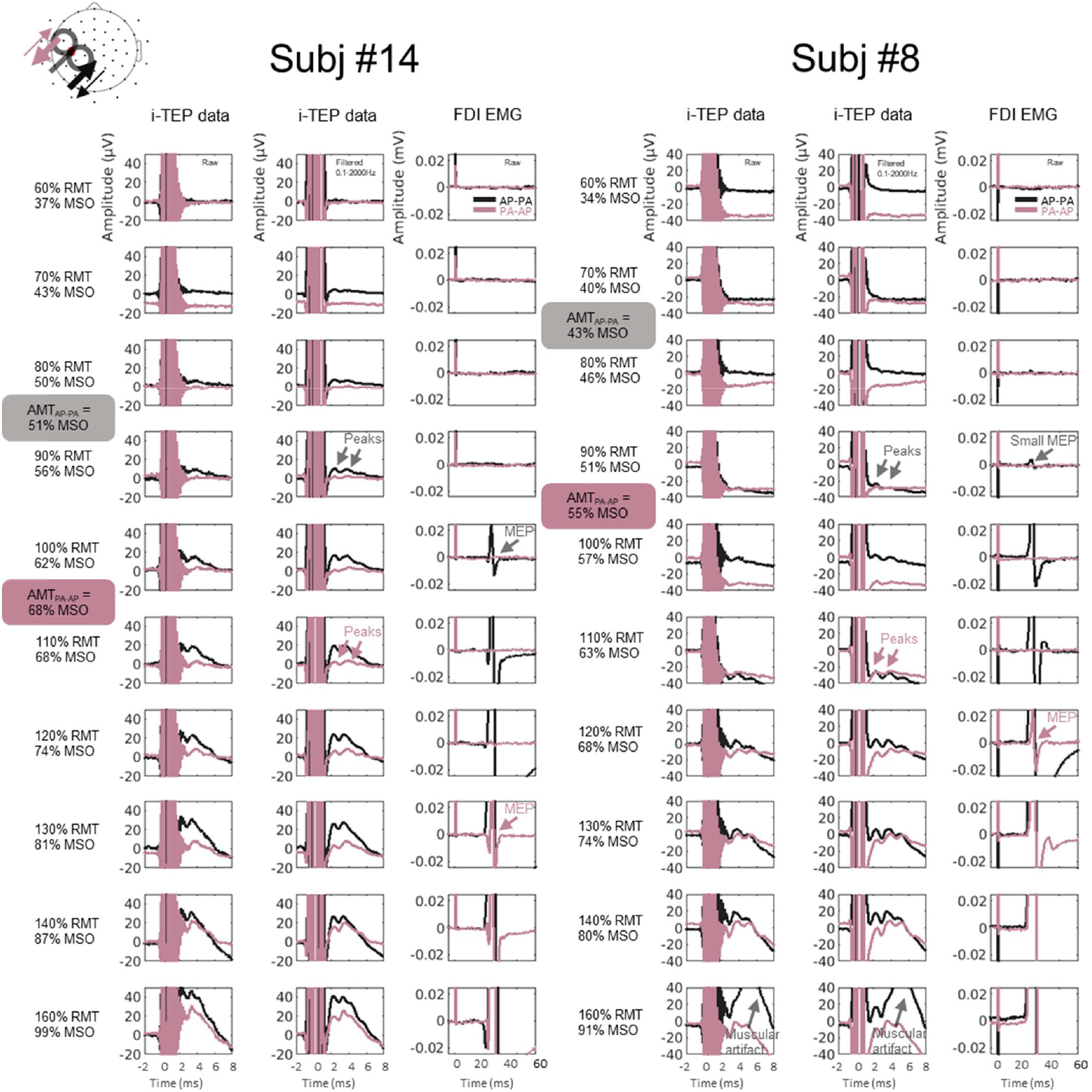
i-TEPs and MEPs are both influenced by current direction. i-TEP responses from electrode close to site of stimulation following M1 stimulation at different intensities (relative to RMT_AP-PA_) and with different current directions (AP-PA vs. PA-AP) for two subjects (for remaining two subjects, see Figure S9). There is a correspondence between the emergence of i-TEPs and MEPs for the specific current directions. i-TEPs and MEPs emerge earlier for AP-PA stimulation compared to PA-AP stimulation. i-TEPs generally emerge at peri-threshold intensities.

## Discussion

By minimizing pulse and muscular twitch artifacts, we were able to detect very early EEG responses to single-pulse TMS of left M1_HAND_ in healthy volunteers. These i-TEPs started approximately 2 ms after the stimulation and consisted of a series of 2-4 higher-frequency peaks with inter-peak intervals of 1.1-1.4 ms superimposed on a slower positive potential lasting up to 6 ms. The i-TEPs were greatest in electrodes overlying the stimulated pericentral cortex and only appeared when TMS was targeting the M1_HAND_ but not the control areas. The i-TEPs were modulated by stimulation intensity and the direction of the induced current. In the following, we discuss the properties of the i-TEPs and how they may corroborate a cortical origin of the i-TEPs. We then touch on potential methodological confounds and how they may have affected the i-TEP recordings.

### Consistency and variability

Single-pulse TMS of M1_HAND_ consistently evoked i-TEPs in all participants when stimulating with peri-threshold intensities (Figure 1B). Moreover, i-TEPs displayed small between-subject variability in latencies (Figure 1C), which could be explained by the fact that TMS leads to a highly synchronized excitation of many pyramidal neurons in the crown of the pericentral gyri. In fact, the i-TEP latency distribution resembles that of the early cortical EEG peak following deep electrical stimulation of the ventral intermediate thalamic nucleus [19], which projects directly to the cortex resulting in highly consistent time locking of cortical potentials.

### Dose-response relationship

The i-TEP peaks were consistently greater in amplitude and more peaks emerged at higher stimulation intensities. This pattern is comparable to the intensity-response relationships found with epidural spinal recordings for TMS-evoked descending corticospinal volleys [20–22]. However, one also needs to consider that the artifact also may increase with stimulus intensity, and thus, may alter the appearance of the i-TEP in an intensity-dependent fashion. Due to the 5000 Hz ringing artifact caused by the TMS pulse, it was also necessary to apply a band-pass filter to the recorded EEG data to estimate the amplitude and latency of the first immediate peak. This did not seem to distort the data features of interest (Figure S1). Yet, caution is still warranted when drawing conclusions from extracted i-TEP peak values, as it is uncertain how the peaks are affected by the decay artifact.

### Site-specificity

While i-TEPs were consistently evoked across subjects when targeting M1_HAND,_ we did not observe any i-TEP-*like* activity when stimulating medial or lateral parietal sites in any individuals, not even when increasing stimulation intensity to match the skull to cortex distance at the site of stimulation (Figure S6). The observed i-TEPs following M1_HAND_ stimulation were composed by high-frequency activity superimposed on a slower positive deflection lasting from approx. 2 to 6 ms. The topographies of the peaks and troughs of the high-frequency activity support the notion that the slower positive deflection is generated in the stimulated pericentral cortex. This is in line with the very first report of TEP topography following M1_HAND_ stimulation, showing a strong activation in ipsilateral motor areas approx. 3 ms after TMS [1]. At the same time, the topographies generated by subtracting the peaks from the troughs of the multi-peak high-frequency response suggest that also this component of the i-TEP might originate from a precentral source. The phenomenon of a high frequency oscillation superimposed on a slower response has previously been observed in EEG recordings. For example, for somatosensory evoked potentials (SSEPs) elicited by electrical stimulation of the median nerve, the slower negative wave observed at around 20ms after stimulation (N20) is combined with high frequency oscillations (>600Hz) generated in the same cortical region [23,24]. Furthermore, results from our lab have confirmed and elaborated this notion by linking high-frequency somatosensory responses to high-frequency changes in intracortical excitability of M1_HAND_ probed with a short-latency intracortical facilitation (SICF) paired-pulse protocol [25]. Furthermore, comparable cortical responses can be recorded from the motor cortex following deep electrical stimulation of the subthalamic nucleus likely via antidromic activation of the hyperdirect pathway [26]. In the light of this body of work, the site specificity of the patterned high-frequency i-TEP response strengthens the indication that they are a pericentral phenomenon. Indeed, the precentral gyrus contains circuitries capable of generating high frequency activity (>600Hz) in response to TMS, as revealed from both invasive epidural recordings at the level of the spinal cord following a single TMS pulse [27] and non-invasively with SICF protocols [28,29]. There are also other possible explanations for the inability to elicit i-TEPs outside of M1_HAND_ in two control sites: Though our stimulation of posterior lateral and midline targets likely affected both gyral crowns and lips granted the relatively large electrical field induced by the 70 mm coil in these conditions, we did not systematically optimize stimulation procedures (e.g., orientation of the coil) based on evoked central or peripheral responses, as we did for M1_HAND_ [15]. It is therefore not possible for us to exclude that the lack of i-TEPs in these parietal regions reflects suboptimal activation of the underlying gray matter in the targeted control regions.

### Impact of induced current direction

Changing the TMS current direction inverted the polarity of the TMS pulse artefact, but not the polarity of the i-TEPs. This finding mirrors the behavior of subdural evoked cortical potentials [26]. These potentials also display inverted stimulation artifacts after reversal of the electrical current during deep brain stimulation of the hyper-direct pathway, while this is not the case for the antidromically evoked cortical activity [26]. Changing the current direction from AP-PA to PA-AP while maintaining the absolute stimulation intensity optimized for AP-PA stimulation resulted in diminished or abolished i-TEPs. Similar patterns are observed for MEPs, suggesting that PA-AP currents are less effective in exciting the pyramidal tract neurons that generate the descending activity in the corticospinal tract. Furthermore, when stimulating at 110% of the RMT for the respective current directions, it was often observed that the amplitudes of Peak 1 were larger following AP-PA stimulation, while the amplitudes of Peak 2 and Peak 3 were greater following PA-AP stimulation (Figure 4DE). This shift in weighting towards the later peaks is also in agreement with recordings of descending waves in the corticospinal tract at the level of the cervical spinal cord when reversing the biphasic pulse [12]. Collectively, this shows that PA-AP waveforms generated i-TEPs and MEPs at higher absolute stimulation intensities and with different qualitative and quantitative properties, resembling features previously demonstrated in epidural recordings of descending corticospinal activity and MEPs [30,31]. This points towards a physiological relationship between i-TEPs and corticospinal activity. However, it should be mentioned that, given the opposing polarities of the stimulation artifact, the two current directions induced somewhat different decay artifacts in the first milliseconds after stimulation. This might have affected the estimation of the amplitude of the peaks. Furthermore, comparisons between phasic activity in EEG and epidural recordings from the corticospinal tract should be made with great caution as other sources than precentral projection neurons may contribute to the EEG signal (e.g., postcentral pyramidal neurons).

### Methodological considerations

Previous TEP research has focused on peaks occurring at latencies later than 10 ms after the TMS pulse [32], because the immediate responses are typically confounded by physiological and instrument artefacts. The main physiological source that obscures the immediate brain responses is caused by peripheral co-stimulation of motor nerve fibers by the TMS-induced electrical field, inducing compound muscle potentials and contractions in neighboring cranial muscles. These direct muscle responses to TMS over M1_HAND_ are seen after ∼5-8 ms, are magnitudes larger than typical EEG responses, and appear as a dipolar pattern over the temporalis muscle [5]. We believe that i-TEPs are unlikely to reflect evoked activity in scalp muscles because the i-TEP appears close to the M1_HAND_ stimulation site, has shorter latencies (first peak after ∼2 ms), and smaller amplitudes than typical muscle artefacts [5]. Furthermore, we took several precautions during data acquisition to avoid scalp muscle activation, including visualizing EEG responses to TMS online as data was collected, and optimizing the coil position by tilting and rotating the coil if needed [5,15]. The TMS-induced electrical field also causes somatosensory and auditory stimulation that evoke peripherally generated EEG responses [3]. It is unlikely that auditory stimulation contributed substantially to the reported i-TEPs, since we used a masking procedure to mask or minimize the click produced during TMS, and because we would expect comparable responses across stimulation sites, which was not the case when stimulating posterior control regions. Somatosensory input from the scalp is transferred to the brain via the trigeminal nerve. Although cortical responses to trigeminal nerve stimulation have been reported already after a few milliseconds [33,34] they show widespread topographies and have shorter inter-peak intervals (<1 ms) [33,34], or have slower onsets (earliest ∼7 ms) and are largest contralateral to stimulation [35–37]. In contrast, we observed larger amplitude responses lateralized to the stimulated hemisphere at latencies below 5 ms and with interpeak intervals of ∼1.1-1.4ms, making trigeminal contamination of the i-TEPs unlikely.

Electromagnetic or vibratory artifacts constitute another potentially confounding source [8,38,39]. However, such artifacts should also have been present after stimulation of the two parietal control sites. As the i-TEP was only elicited by M1_HAND_ stimulation, we consider it unlikely that it represents an electromagnetic or vibratory artifact. Furthermore, when the current of the biphasic pulse was reversed, the pulse artifact reversed its polarity, but not the polarity of the i-TEP peaks.

It should be noted that a previous attempt to unveil the immediate response to TMS found EEG responses with similar frequencies to the ones observed here [40]. However, these responses were constrained to a single electrode underneath the stimulation coil, did not depend on the stimulation site, and were also observed when stimulating a phantom (melon) [40], implying a non-physiological origin. Given these features, it was, in that study, proposed that the TMS discharge induced vibration of the EEG cables through either physical contact, airborne sound waves, or electromagnetic interactions [40]. In contrast, the i-TEPs reported here had a more widespread (lateralized) topography and were absent when stimulating other areas of the scalp or a phantom (knee) using similar intensities as when targeting the M1_HAND_. Taken together, these results support the notion that the observed i-TEPs were generated in the stimulated M1_HAND_ rather than having an artifactual origin. However, future studies on the i-TEP still need to pay close attention to the potential contribution of physiological and instrumental artifacts.

We would also like to alert the reader to some general limitations of the present study. First, we tested a relatively small number of young healthy individuals. Although we were able to consistently evoke i-TEPs in all participants, the ability to reliably evoke i-TEPs with single-pulse TMS of M1_HAND_ needs to be replicated in larger cohorts. Second, we used biphasic stimulation that is less direction selective than monophasic pulses in recruiting cortical neurons [4]. We chose a biphasic pulse waveform, because they show shorter decay artifacts following stimulation [41] and elicit MEPs at lower stimulation intensities [13], which lowered the risk of electrical co-stimulation of neighboring scalp muscles. However, the use of biphasic pulse waveforms compared to monophasic pulses may have masked some of the direction-specific differences in i-TEP phenomenology between AP and PA dominant currents in a similar manner as has been reported for epidurally recorded I-waves [42]. That said, recent work has shown that also for biphasic pulses, PA-AP and AP-PA directed currents excite different neuronal subpopulations in the pericentral cortex resulting in different patterns of descending corticospinal volleys [27,43,44].

## Conclusion

By minimizing physiological and instrumental artifacts, we observed an immediate EEG response to TMS across all tested subjects. These responses were dependent on stimulation intensity and current direction and were only present when stimulating M1_HAND_. Given the rhythmicity of around 650-800 Hz and the topographical distribution of the potentials of i-TEPs in the Rolandic area, our results provide a clear testable hypothesis for future research, namely that i-TEPs are a direct marker of TMS-induced excitation of M1_HAND_, and the cortical source of the indirect corticospinal volleys (I-waves) that have been demonstrated in invasive epidural recordings. We hope that our report will stimulate research into the cortical origin of i-TEPs and may turn out to be a useful clinical marker to probe M1_HAND_ excitability in patients with motor disorders.

## Acknowledgements

This work was supported by a Grand Solutions grant “Precision Brain-Circuit Therapy - Precision-BCT” from Innovation Fund Denmark to Hartwig R. Siebner (grant no. 9068-00025B) and a Collaborative Project grant “ADAptive and Precise Targeting of cortex-basal ganglia circuits in Parkinson’s Disease - ADAPT-PD” from The Lundbeck Foundation to Hartwig R. Siebner (grant no. R336-2020-1035). Mikkel M. Beck is funded by a post doc grant from the Capital Region Denmark (Region H). Lasse Christiansen holds a postdoc grant from The Lundbeck Foundation (grant no. R322-2019-2406). Leo Tomasevic holds an ‘Experiment grant’ from The Lundbeck Foundation (grant no. R346-2020-1822). Axel Thielscher was supported by The Lundbeck Foundation (grant no. R313-2019-622).

## Author Contributions

Conceptualization: MMB, LC, LT, HRS; Data curation: MMB, MAJM, LT; Formal analysis: MMB, LC, LT; Funding acquisition: HRS; Investigation: MMB, LC, MAJM, AJF, LT; Methodology: MMB, LC, MAJM, AJF, MCV, AT, TOB, LT, HRS; Project Administration: MMB; Resources: HRS; Visualization: MMB, LC, LT, HRS; Writing – original draft: MMB, LC, HRS, LT; Writing – review & editing: MMB, LC, MAJM, AJF, MCV, AT, TOB, LT, HRS

## References

[1] Ilmoniemi RJ, Virtanen J, Ruohonen J, Karhu J, Aronen HJ, Näätänen R, et al. Neuronal responses to magnetic stimulation reveal cortical reactivity and connectivity. Neuroreport 1997;8:3537–40. 10.1097/00001756-199711100-00024.

[2] Ilmoniemi RJ, Kičić D. Methodology for combined TMS and EEG. Brain Topogr 2010;22:233–48. 10.1007/s10548-009-0123-4.

[3] Conde V, Tomasevic L, Akopian I, Stanek K, Saturnino GB, Thielscher A, et al. The non-transcranial TMS-evoked potential is an inherent source of ambiguity in TMS-EEG studies. Neuroimage 2019;185:300–12. 10.1016/J.NEUROIMAGE.2018.10.052.

[4] Siebner HR, Funke K, Aberra AS, Antal A, Bestmann S, Chen R, et al. Transcranial magnetic stimulation of the brain: What is stimulated? – A consensus and critical position paper. Clin Neurophysiol 2022;140:59–97. 10.1016/J.CLINPH.2022.04.022.

[5] Mutanen T, Mäki H, Ilmoniemi RJ. The effect of stimulus parameters on {TMS}-{EEG} muscle artifacts. Brain Stimul 2013;6:371–6. 10.1016/j.brs.2012.07.005.

[6] Veniero D, Bortoletto M, Miniussi C. TMS-EEG co-registration: On TMS-induced artifact. Clin Neurophysiol 2009;120:1392–9. 10.1016/j.clinph.2009.04.023.

[7] Tomasevic L, Takemi M, Siebner HR. Synchronizing the transcranial magnetic pulse with electroencephalographic recordings effectively reduces inter-trial variability of the pulse artefact. PLoS One 2017;12:e0185154. 10.1371/journal.pone.0185154.

[8] Bergmann TO, Tomasevic L, Siebner HR. Transcranial brain stimulation and EEG/MEG. Oxford Handb. Transcranial Stimul. Second Ed., Oxford University Press; 2021. 10.1093/oxfordhb/9780198832256.013.26.

[9] Patton HD, Amassian VE. Single and multiple-unit analysis of cortical stage of pyramidal tract activation. J Neurophysiol 1954;17:345–63. 10.1152/jn.1954.17.4.345.

[10] Adrian ED, Moruzzi G. Impulses in the pyramidal tract. J Physiol 1939;97:153–99. 10.1113/jphysiol.1939.sp003798.

[11] Di Lazzaro V, Oliviero A, Profice P, Saturno E, Pilato F, Insola A, et al. Comparison of descending volleys evoked by transcranial magnetic and electric stimulation in conscious humans. Electroencephalogr Clin Neurophysiol -Electromyogr Mot Control 1998;109:397–401. 10.1016/S0924-980X(98)00038-1.

[12] Lazzaro V Di, Oliviero A, Profice P, Meglio M, Cioni B, Tonali P, et al. Descending spinal cord volleys evoked by transcranial magnetic and electrical stimulation of the motor cortex leg area in conscious humans. J Physiol 2001;537:1047–58. 10.1111/j.1469-7793.2001.01047.x.

[13] Kammer T, Beck S, Thielscher A, Laubis-Herrmann U, Topka H. Motor thresholds in humans: A transcranial magnetic stimulation study comparing different pulse waveforms, current directions and stimulator types. Clin Neurophysiol 2001;112:250–8. 10.1016/S1388-2457(00)00513-7.

[14] Yousry TA, Schmid UD, Alkadhi H, Schmidt D, Peraud A, Buettner A, et al. Localization of the motor hand area to a knob on the precentral gyrus. A new landmark. Brain 1997;120:141–57. 10.1093/brain/120.1.141.

[15] Casarotto S, Fecchio M, Rosanova M, Varone G, D’Ambrosio S, Sarasso S, et al. The rt-TEP tool: real-time visualization of TMS-Evoked Potentials to maximize cortical activation and minimize artifacts. J Neurosci Methods 2022;370:109486. 10.1016/J.JNEUMETH.2022.109486.

[16] Dockès J, Poldrack RA, Primet R, Gözükan H, Yarkoni T, Suchanek F, et al. Neuroquery, comprehensive meta-analysis of human brain mapping. Elife 2020;9. 10.7554/ELIFE.53385.

[17] Stokes MG, Chambers CD, Gould IC, English T, McNaught E, McDonald O, et al. Distanceadjusted motor threshold for transcranial magnetic stimulation. Clin Neurophysiol 2007;118:1617–25. 10.1016/j.clinph.2007.04.004.

[18] Russo S, Sarasso S, Puglisi GE, Dal PalùD, Pigorini A, Casarotto S, et al. TAAC -TMS Adaptable Auditory Control: A universal tool to mask TMS clicks. J Neurosci Methods 2022;370:109491. 10.1016/J.JNEUMETH.2022.109491.

[19] Walker HC, Huang H, Gonzalez CL, Bryant JE, Killen J, Knowlton RC, et al. Short latency activation of cortex by clinically effective thalamic brain stimulation for tremor. Mov Disord 2012;27:1404–12. 10.1002/MDS.25137.

[20] Burke D, Hicks R, Gandevia SC, Stephen J, Woodforth I, Crawford M. Direct comparison of corticospinal volleys in human subjects to transcranial magnetic and electrical stimulation. J Physiol 1993;470:383–93. 10.1113/JPHYSIOL.1993.SP019864.

[21] Kaneko K, Kawai S, Fuchigami Y, Shiraishi G, Ito T. Effect of stimulus intensity and voluntary contraction on corticospinal potentials following transcranial magnetic stimulation. J Neurol Sci 1996;139:13–4.

[22] Di Lazzaro V, Oliviero A, Profice P, Insola A, Mazzone P, Tonali P, et al. Effects of voluntary contraction on descending volleys evoked by transcranial electrical stimulation over the motor cortex hand area in conscious humans. Exp Brain Res 1999;124:525–8. 10.1007/s002210050649.

[23] Curio G, Mackert BM, Burghoff M, Koetitz R, Abraham-Fuchs K, Härer W. Localization of evoked neuromagnetic 600 Hz activity in the cerebral somatosensory system. Electroencephalogr Clin Neurophysiol 1994;91:483–7. 10.1016/0013-4694(94)90169-4.

[24] Hashimoto I, Mashiko T, Imada T. Somatic evoked high-frequency magnetic oscillations reflect activity of inhibitory interneurons in the human somatosensory cortex. Electroencephalogr Clin Neurophysiol -Evoked Potentials 1996;100:189–203. 10.1016/0168-5597(95)00244-8.

[25] Tomasevic L, Siebner R, Thielscher A, Manganelli F, Pontillo G, Dubbioso R. Relationship between high-frequency activity in the cortical sensory and the motor hand areas, and their myelin content 2022. 10.1016/j.brs.2022.04.018.

[26] Miocinovic S, de Hemptinne C, Chen W, Isbaine F, Willie JT, Ostrem JL, et al. Cortical potentials evoked by subthalamic stimulation demonstrate a short latency hyperdirect pathway in humans. J Neurosci 2018;38:9129–41. 10.1523/JNEUROSCI.1327-18.2018.

[27] Di Lazzaro V, Rothwell JC. Corticospinal activity evoked and modulated by non-invasive stimulation of the intact human motor cortex. J Physiol 2014;592:4115–28. 10.1113/jphysiol.2014.274316.

[28] Tokimura H, Ridding MC, Tokimura Y, Amassian VE, Rothwell JC. Short latency facilitation between pairs of threshold magnetic stimuli applied to human motor cortex. Electroencephalogr Clin Neurophysiol -Electromyogr Mot Control 1996;101:263–72. 10.1016/0924-980X(96)95664-7.

[29] Ziemann U, Tergau F, Wassermann EM, Wischer S, Hildebrandt J, Paulus W. Demonstration of facilitatory I wave interaction in the human motor cortex by paired transcranial magnetic stimulation. J Physiol 1998;511:181–90. 10.1111/j.1469-7793.1998.181bi.x.

[30] Mills KR, Boniface SJ, Schubert M. Magnetic brain stimulation with a double coil: the importance of coil orientation. Electroencephalogr Clin Neurophysiol Potentials Sect 1992;85:17–21. 10.1016/0168-5597(92)90096-T.

[31] Brasil-Neto JP, Cohen LG, Panizza M, Nilsson J, Roth BJ, Hallet M. Optimal focal transcranial magnetic activation of the human motor cortex: effects of coil orientation, shape of the induced current pulse, and stimulus intensity. J Clin Neurophysiol 1992;9:132–6.

[32] Farzan F, Bortoletto M. Identification and verification of a “true” TMS evoked potential in TMS-EEG. J Neurosci Methods 2022;378:109651. 10.1016/J.JNEUMETH.2022.109651.

[33] Leandri M, Parodi CI, Favale E. Early evoked potentials detected from the scalp of man following infraorbital nerve stimulation. Electroencephalogr Clin Neurophysiol Evoked Potentials 1985;62:99–107. 10.1016/0168-5597(85)90021-8.

[34] Leandri M, Parodi CI, Zattoni J, Favale E. Subcortical and cortical responses following infraorbital nerve stimulation in man. Electroencephalogr Clin Neurophysiol 1987;66:253–62. 10.1016/0013-4694(87)90074-5.

[35] Hashimoto I. Trigeminal evoked potentials following brief air puff: Enhanced signal‐to‐noise ratio. Ann Neurol 1988;23:332–8. 10.1002/ana.410230404.

[36] Nevalainen P, Ramstad R, Isotalo E, Haapanen ML, Lauronen L. Trigeminal somatosensory evoked magnetic fields to tactile stimulation. Clin Neurophysiol 2006;117:2007–15. 10.1016/j.clinph.2006.05.019.

[37] Findler G, Feinsod M. Sensory evoked response to electrical stimulation of the trigeminal nerve in humans. J Neurosurg 1982;56:545–9. 10.3171/jns.1982.56.4.0545.

[38] Li B, Virtanen JP, Oeltermann A, Schwarz C, Giese MA, Ziemann U, et al. Lifting the veil on the dynamics of neuronal activities evoked by transcranial magnetic stimulation. Elife 2017;6. 10.7554/elife.30552.

[39] Mueller JK, Grigsby EM, Prevosto V, Petraglia FW, Rao H, Deng Z-D, et al. Simultaneous transcranial magnetic stimulation and single-neuron recording in alert non-human primates 2014. 10.1038/nn.3751.

[40] Pillen S, Knodel N, Zrenner C, Ziemann U, Bergmann T. A setup for studying very early TMS-evoked EEG potentials: prospects and pitfalls. Brain Stimul 2019;12:565–6. 10.1016/j.brs.2018.12.873.

[41] Rogasch NC, Thomson RH, Daskalakis ZJ, Fitzgerald PB. Short-latency artifacts associated with concurrent TMS-EEG. Brain Stimul 2013;6:868–76. 10.1016/j.brs.2013.04.004.

[42] Di Lazzaro V, Oliviero A, Mazzone P, Insola A, Pilato F, Saturno E, et al. Comparison of descending volleys evoked by monophasic and biphasic magnetic stimulation of the motor cortex in conscious humans. Exp Brain Res 2001;141:121–7. 10.1007/s002210100863.

[43] Federico P, Perez MA. Distinct Corticocortical Contributions to Human Precision and Power Grip. Cereb Cortex 2017;27:5070–82. 10.1093/cercor/bhw291.

[44] Hamada M, Galea JM, Di Lazzaro V, Mazzone P, Ziemann U, Rothwell JC. Two Distinct Interneuron Circuits in Human Motor Cortex Are Linked to Different Subsets of Physiological and Behavioral Plasticity. J Neurosci 2014;34:12837–49. 10.1523/JNEUROSCI.1960-14.2014.

