## Supplementary figures and tables for "Transcranial magnetic stimulation of primary motor cortex elicits a site-specific immediate transcranial evoked potential"

*Shared authorship

#Correspodance to:

**Leo Tomasevic**
Danish Research Centre for Magnetic Resonance (DRCMR)
Kettegård Allé 30, 2650 Hvidovre, Denmark


### Supplementary results

#### Overview of experimental conditions

Table S1 provides an overview of the conditions and the number of trials performed and included for each subject from the main- and control experiments.

**Table S1|Overview of experimental conditions and number of trials**

| | M1 | | | | | PPC | | Midline |
|---|---|---|---|---|---|---|---|---|
| | AP-PA 95% $RMT_{AP-PA}$ | AP-PA 110% $RMT_{AP-PA}$ | PA-AP 110% $RMT_{AP-PA}$ | PA-AP 110% $RMT_{PA-AP}$ | Input-output | AP-PA 110% $RMT_{AP-PA}$ | Scalp-to-cortex adjusted | AP-PA 110% $RMT_{AP-PA}$ |
| S1 | 100 | 100 | 100 | 99 | 50 | 100 | 100 | 100 |
| S2 | 100 | 100 | 100 | 100 | % | 100 | % | % |
| S3 | 65 | 100 | % | % | % | % | % | 100 |
| S4 | 98 | 100 | 100 | 97 | % | 99 | % | 100 |
| S5 | 97 | 98 | 100 | 92 | % | 98 | % | 97 |
| S6 | 100 | 96 | 94 | 100 | % | 90 | % | 90 |
| S7 | 100 | 100 | 100 | 100 | % | 100 | 98 | 99 |
| S8 | 100 | 96 | 99 | 99 | 50 | 96 | 100 | 98 |
| S9 | 96 | 98 | 97 | % | % | 69 | % | 97 |
| S10 | 100 | 100 | % | % | % | % | % | % |
| S11 | 96 | 100 | % | % | % | % | % | 100 |
| S12 | 98 | 99 | % | % | % | % | % | % |
| S13 | % | % | % | % | 50 | % | % | % |
| S14 | % | % | % | % | 50 | % | % | % |
| Avg | 95,8 | 98.9 | 98,8 | 98,1 | 50 | 94.0 | 99.3 | 97.9 |
| SD | 9.8 | 1.6 | 2,2 | 2,9 | 0 | 10.7 | 1.2 | 3.2 |

Table S1 displays the number of trials included for averaging for each subject. '%' reflects that the given condition was not performed for that subject.

#### Stimulation parameters

Table S2 provides an overview of participants' RMT, extracted MEP amplitudes, and latencies following M1 stimulations for the different stimulation conditions in the main experiment.

**Table S2| Stimulation parameters, MEP peak-to-peak amplitudes and latencies.**

| | RMT | | Peak-to-peak amplitudes of MEPs (mV) | | | | Latencies of MEPs (ms) | | | |
|---|---|---|---|---|---|---|---|---|---|---|
| | AP-PA | PA-AP | AP-PA 95% $RMT_{AP-PA}$ | AP-PA 110% $RMT_{AP-PA}$ | PA-AP 110% $RMT_{AP-PA}$ | PA-AP 110% $RMT_{PA-AP}$ | AP-PA 95% RMT $_{AP-PA}$ | AP-PA 110% $RMT_{AP-PA}$ | PA-AP 110% $RMT_{AP-PA}$ | PA-AP 110% $RMT_{PA-AP}$ |
| S1 | 35 | 44 | 0.05±0.06 | 0.70±0.73 | 0.03±0.02 | 0.09 ± 0.08 | 24.8 | 25.2±1.3 | - | 25.7±2.6 |
| S2 | 40 | 55 | 0.10±0.20 | 0.44±0.61 | 0.02±0.01 | 0.87 ± 0.69 | 24.4 | 23.4±1.3 | - | 23.7±0.6 |
| S3 | 40 | % | 0.04±0.08 | 2.33±2.51 | % | % | - | 24.6±4.5 | % | % |
| S4 | 35 | 40 | 0.02±0.01 | 1.14±0.86 | 0.06±0.07 | 0.08 ± 0.07 | - | 23.2±0.9 | 25.4 | 27.3±7.8 |
| S5 | 31 | 38 | 0.11±0.21 | 0.65±0.81 | 0.06±0.14 | 3.19 ± 2.40 | 25.8 | 24.9±1.1 | 26.8 | 25.6±0.7 |
| S6 | 43 | | 0.08±0.09 | 0.25±0.31 | 0.05±0.04 | 0.26 ± 0.30 | 24.6 | 24.9±1.1 | 25.5 | 25.4±1.9 |
| S7 | 32 | 38 | 0.02±0.02 | 0.23±0.37 | 0.08±0.09 | 0.19 ± 0.19 | - | 23.4±1.1 | 25.4 | 24.6±1.5 |
| S8 | 31 | 39 | 0.07±0.19 | 1.15±0.93 | 0.10±0.26 | 3.13 ± 1.64 | 21.7 | 22.9±6.7 | 26.6 | 25.5±1.3 |
| S9 | 47 | 62 | 0.05±0.06 | 0.11±0.25 | 0.03±0.01 | % | - | - | - | - |
| S10 | 38 | % | 0.02±0.02 | 0.31±0.34 | % | % | - | 25.6±1.8 | % | % |
| S11 | 30 | % | 0.03±0.03 | 1.06±0.69 | % | % | - | 23.8±1.4 | % | % |
| S12 | 38 | % | 0.02±0.01 | 0.18±0.29 | % | % | - | 26.8±2.9 | % | % |
| Average | 36,7 | 45,1 | 0.05 | 0.71 | 0.05 | 1.12 | 24.1 | 24.4 | 25.9 | 25.4 |
| SD | 5,3 | 9,6 | 0.03 | 0.64 | 0.03 | 1.42 | 1.5 | 2.0 | 0.7 | 1.1 |

Data reported as means ± standard deviations. – no clear MEP response for the given condition; % given condition not performed for the subject.

**Effects of filtering i-TEP data**

In figure S1, we show the effects of band-pass filtering (0.1-2000 Hz) the i-TEP waveform data to remove the ringing from the TMS pulse artefact (5000Hz) that lasts approx. 2.0-2.5ms and rides on top of the first high-frequency i-TEP peak (Peak 1).

(Insert Figure S1 here)

**TMS over M1-HAND elicits prototypical TEP and an i-TEP**

In Figure S2 below, it can be seen how TMS over M1-HAND elicits an EEG waveform response with typical, well-characterized components, including N15, P30, N45, P60, N100 and P180.

(Insert Figure S2 here)

**Effects of varying stimulation intensities over M1 on i-TEPs**

In Figure S3 below, we show the consistent effect of stimulation intensity on immediate TEP responses for all participants following stimulation over M1 and 95% (grey traces) and 110% of $RMT_{AP-PA}$ (black traces) from an electrode close to the stimulation site. For all subjects, peaks were more prominent following suprathreshold stimulation compared to subthreshold stimulation.

(Insert Figure S3 here)

**Effects of varying stimulation site on i-TEPs**

In Figure S4 and S5 below, we show the effect of varying stimulation site on immediate TEP responses for all participants undergoing both M1 and midline stimulation (Figure S4) and M1, Midline and PPC stimulation (Figure S5) at 110% of $RMT_{AP-PA}$. Please note that we have plotted the response from an electrode close to the site of stimulation in all cases as highlighted in bold in the headplots. For all participants, stimulating the M1 at 110% $RMT_{AP-PA}$ resulted in a clear i-TEP response, whereas stimulating the midline or the PPC at the same intensity did not.

(Insert Figure S4 and S5 here)

In Figure S6, we show that adjusting stimulation intensity to account for potential differences in scalp-to-cortex distance did not change the pattern shown in the figures above (Fig. S4-S5). Participants were stimulated with TMS over M1 and PPC. The absolute stimulation intensities (in %MSO) for the RMT-based and the scalp-to-cortex adjusted condition were 39% and 47% for subjects S1, 35% and 45% for subjects S7, and 34% and 51% for subjects S8, respectively. Stimulating the M1 leads to a clear i-TEP, whereas this is not the case for either of the two intensities when targeting the PPC.

(Insert Figure S6 here)

Figure S7 shows responses to 50 TMS pulses delivered over the left knee of a single male subject. Stimulating the knee did not result in any clear responses across nine electrodes following the decay artefact. Stimulation was delivered at an intensity around the average of what was used in the main experiment (40% MSO; AP-PA current direction) with the coil handle oriented perpendicular to the electrode wires (Figure S7). Stimulating over the knee did not elicit any i-TEP-like responses.

(Insert Figure S7 here)

**Impact of varying current direction and intensity over M1 on i-TEPs**

In Figure S8, we show the effect of varying current direction for M1 stimulation on immediate TEP responses for all participants (N=7) undergoing M1 stimulation with biphasic AP-PA and PA-AP stimulation at 110% $RMT_{AP-PA}$ and 110% $RMT_{PA-AP}$. Please refer to supplementary table S2 for the corresponding MEP data. We consistently observed smaller i-TEP responses when stimulating with PA-AP current direction at the same absolute stimulation intensity (110% $RMT_{AP-PA)}$. Increasing the intensity to 110% $RMT_{PA-AP}$ with PA-AP directed currents resulted in an increase in amplitude of the i-TEPs. This was particularly the case for the later i-TEP peaks (Peak 2 and Peak 3).

(Insert Figure S8 here)

Figure S9 presents input-output responses from two additional participants (Subject 1 and 13). The figure displays data from stimulation at intensities ranging from 70-140% $RMT_{AP-PA}$ in steps of 10% $RMT_{AP-PA}$ with both AP-PA and PA-AP current directions. For each subject, the left column displays raw EEG traces from an electrode close to the site of stimulation, the middle column displays filtered (0.1-2000Hz) data from the same electrode while the right column displays corresponding MEP data. The emergence of i-TEPs generally seem to occur later for PA-AP directed currents than for AP-PA currents. This fits with the observation that MEP thresholds are greater for PA-AP stimulation compared to AP-PA stimulation.

(Insert Figure S9 here)

**Impact of re-referencing**

In all previous figures, we have shown raw or filtered data referenced to the forehead as recorded. However, re-referencing to average reference is a common preprocessing step in TMS-EEG studies. In Figure S10 below, we show the grand-averaged i-TEP response for both AP-PA and PA-AP stimulation both using the recorded (forehead) reference and after re-referencing to the average of all electrodes. Re-referencing did not seem to affect the properties of the i-TEPs.

(Insert Figure S10 here)

**Figures and legends**

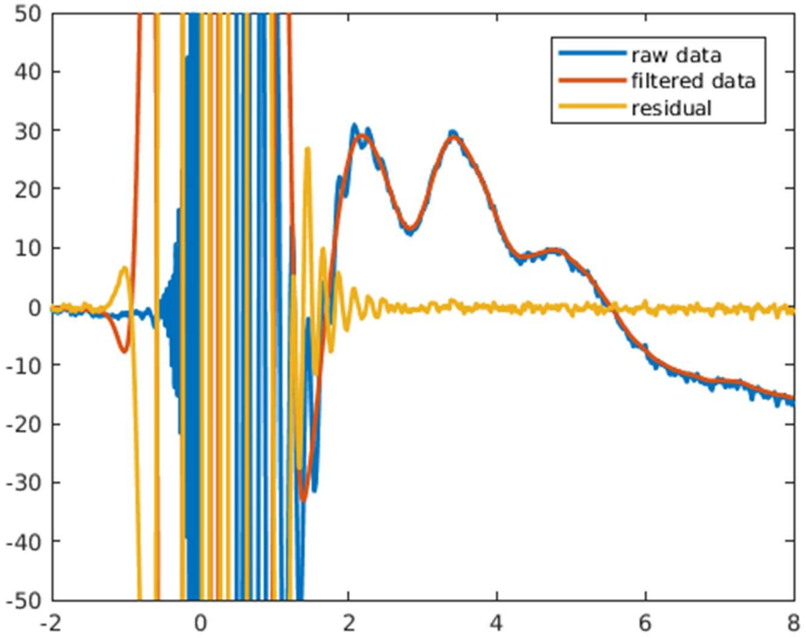


**Figure S1. Effects of filtering on short-latency TMS-evoked potentials in a single representative EEG trace.** Displays raw and filtered (0.1-2000Hz, zero-phase, 2nd order Butterworth) EEG traces in blue and orange, respectively. In yellow, we display the differences between raw and filtered timeseries (residual), highlighting that the filter attenuates the high-frequency (5000Hz) ringing from the TMS pulse lasting ~2.5 ms that rides on top of the first TEP peak.

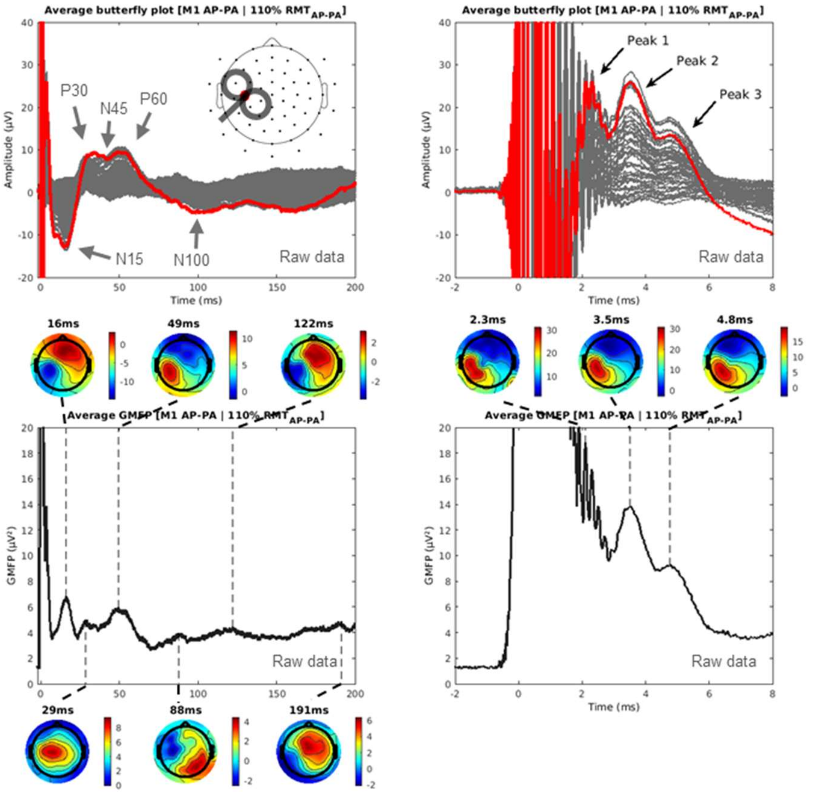


**Figure S2. TMS over M1 elicits prototypical TEP peaks and i-TEPs.** Displays raw group-averaged EEG butterfly plots following biphasic TMS (AP-PA) delivered at 110% $RMT_{AP\text{-}PA}$ over $M1_{HAND}$. Top row displays butterfly plots at two timescales and bottom row the Global Mean Field Power (GMFP)with corresponding topoplots (min-max scaled)(N=12). Trace highlighted in red in top row corresponds to an electrode over left sensorimotor cortex close to the stimulating coil. Suprathreshold stimulation of M1 evokes typical TEP components, including N15, P30, N45, P60 and N100 as seen in left column. Suprathreshold stimulation of M1 also evokes i-TEPs as displayed in right column.

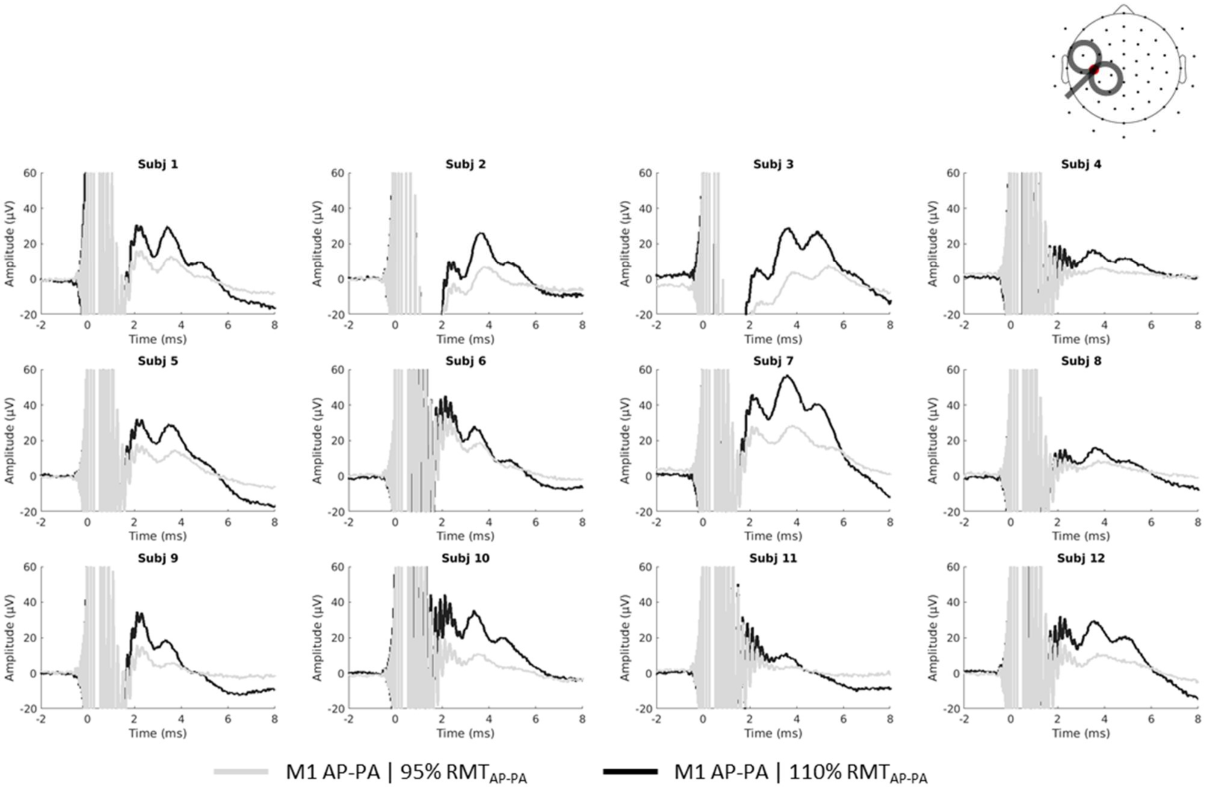


**Figure S3. Effects of stimulation intensity on early TEPs in all tested subjects.** Stimulation over $M1_{HAND}$ at 110% $RMT_{AP-PA}$ (black) or 95% $RMT_{AP-PA}$ (grey) in all tested subjects. Traces are taken from an electrode close to the site of stimulation as shown in the electrode layout plot in the upper right hand corner.

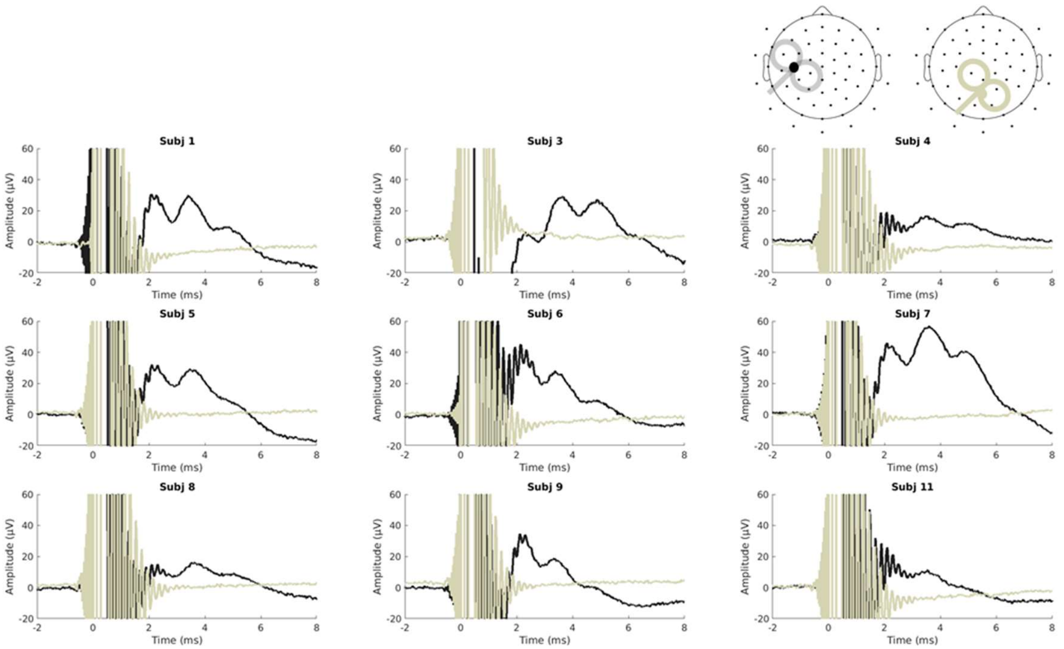

**Figure S4. Effects of stimulation sites on i-TEPs.** Stimulation over M1 (black) or midline (sand) at 110% $RMT_{AP\text{-}PA}$ in all tested subjects. Traces are taken from an electrode close to the site of stimulation as shown in the electrode layout plot.

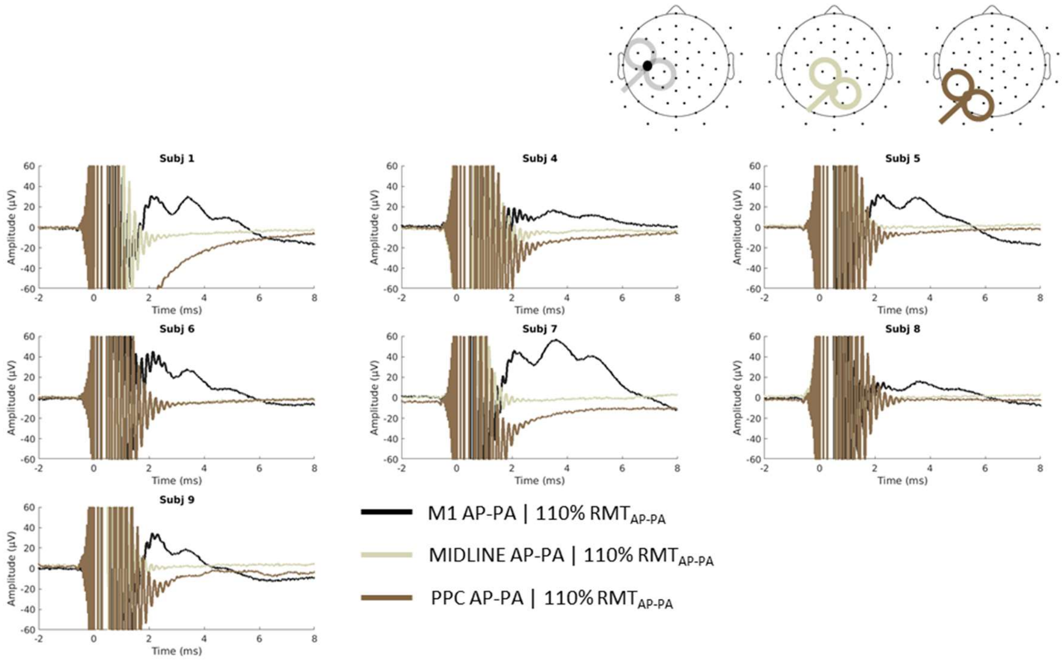


**Figure S5. Effects of stimulation sites on early TEPs.** Stimulation over M1 (black), midline (sand) or PPC (brown) at 110% $RMT_{AP\text{-}PA}$ in all tested subjects. Traces are taken from an electrode close to the site of stimulation as shown in the electrode layout plot.

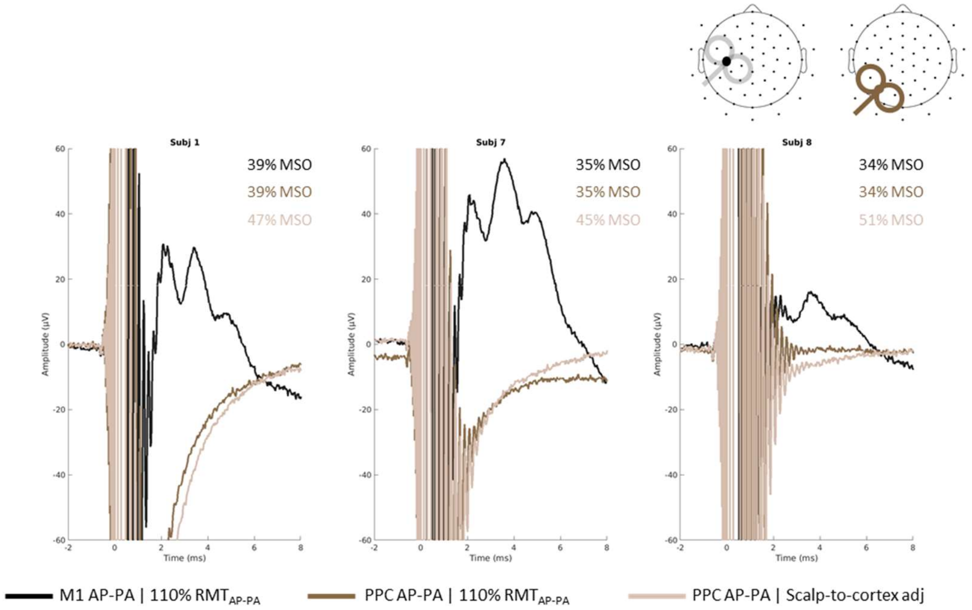


**Figure S6. Effects of stimulation sites on early TEPs.** Plots show early TEPs in all tested subjects receiving stimulation over M1 (black) and over posterior parietal cortex (PPC; brown) at 110% RMT$_{AP-PA}$ (average 36% of MSO) and over PPC with intensity adjusted for differences in scalp-to-cortex distance (average 48% of MSO)(light brown).

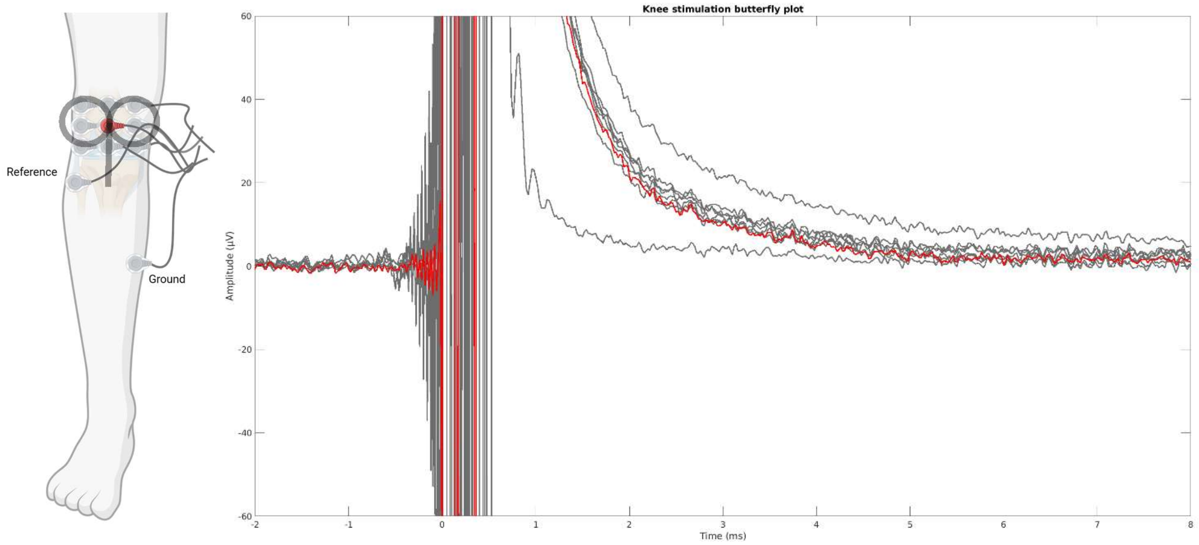


**Figure S7. Responses to TMS of knee.** Responses to TMS delivered at 40% MSO over 9 electrodes positioned on the knee corresponding to the average intensity used for the 110% RMT condition in the main experiment. Impedances were below 5kOhm for all electrodes. The electrode marked in red was closest to site of stimulation.

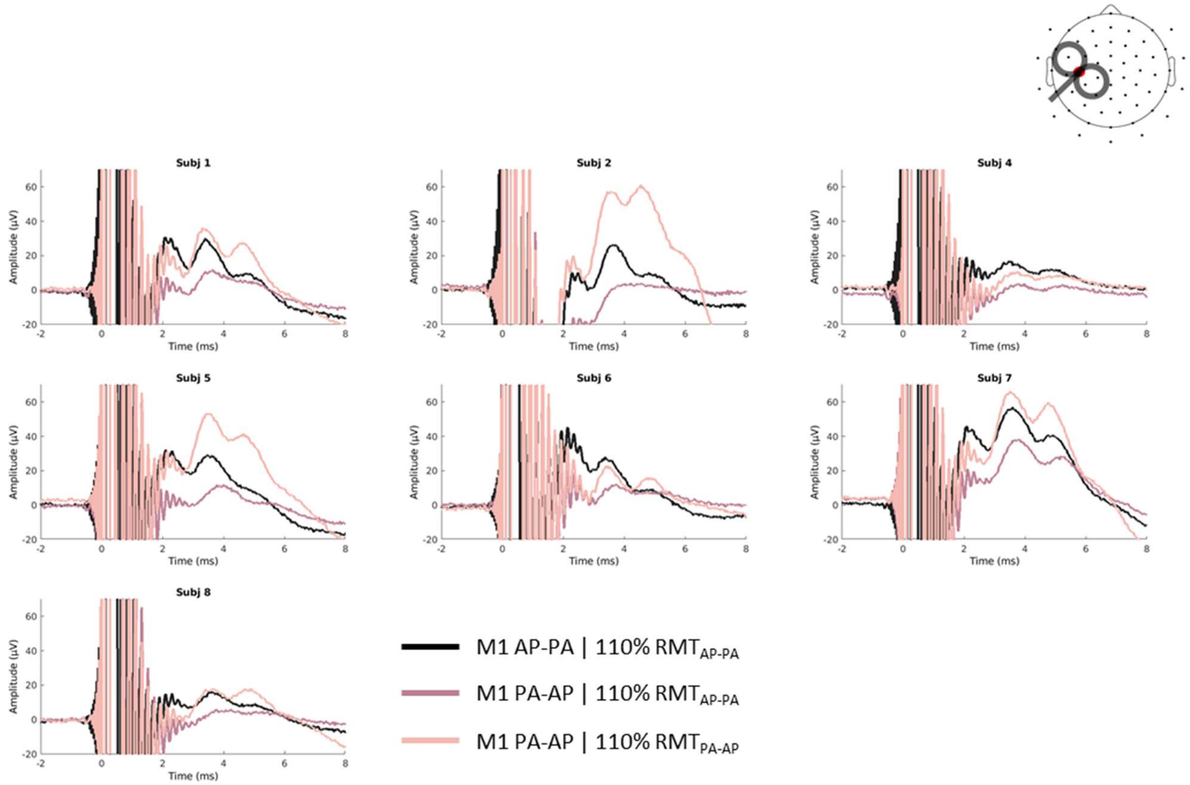


**Figure S8. Effects of current direction on i-TEPs.** i-TEPs in all tested subjects receiving stimulation over M1 with biphasic AP-PA stimulation at 110% RMT$_{AP\text{-}PA}$ (black); biphasic PA-AP stimulation at 110% RMT$_{AP\text{-}PA}$ (pink); and biphasic PA-AP stimulation at 110% RMT$_{PA\text{-}AP}$ (light pink).

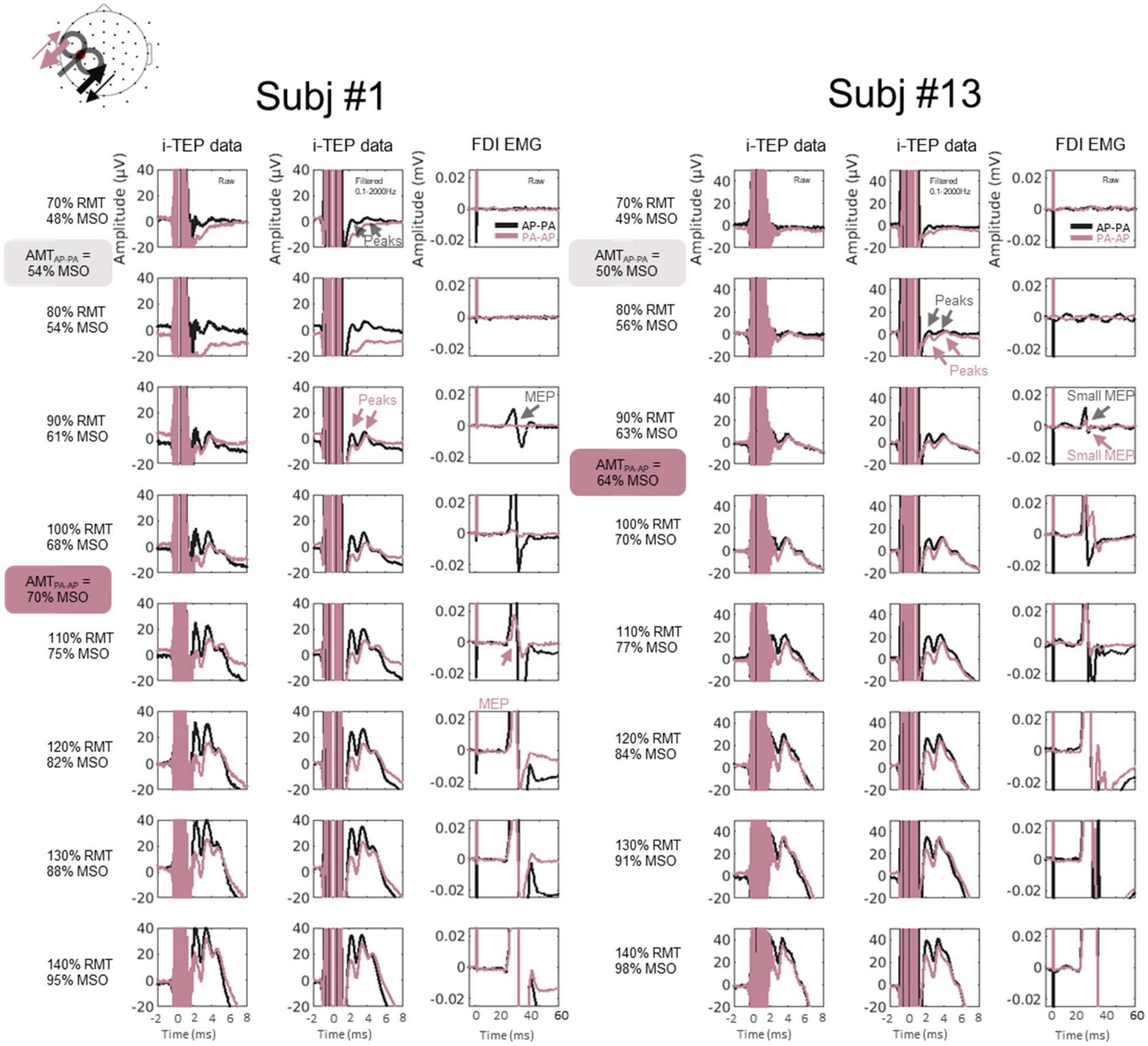


**Figure S9. Effects of stimulation intensity and current direction on i-TEPs and MEPs.** i-TEP responses from electrode close to site of stimulation following M1 stimulation at different intensities (relative to $RMT_{AP-PA}$) and with different current directions (AP-PA vs. PA-AP) for two remaining subjects.

**(A) i-TEP butterfly plots for forehead and average reference for different current directions.**

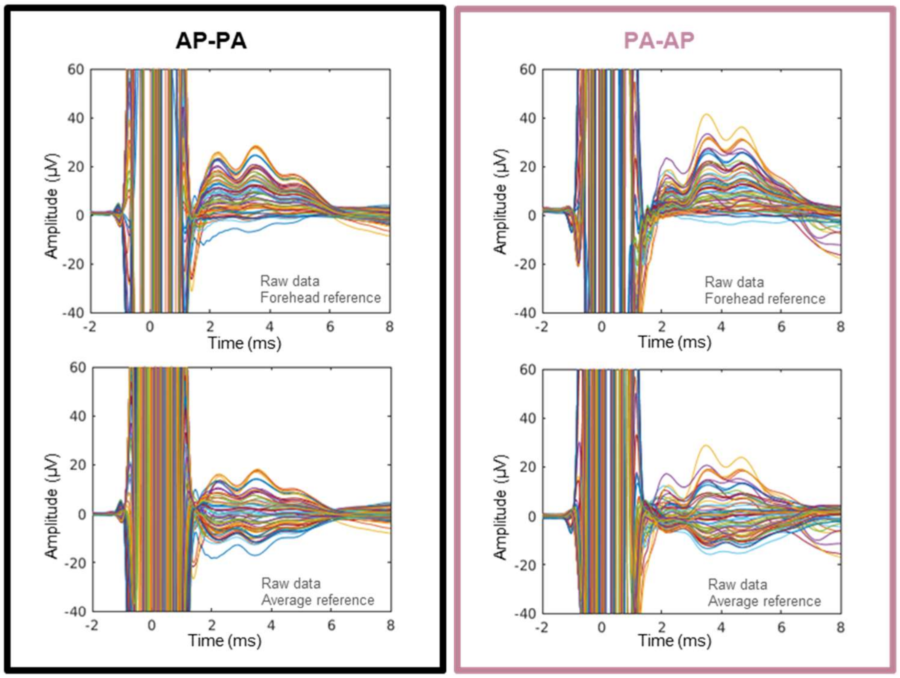


**(B) i-TEP waveforms and topographies for different current directions in average reference.**

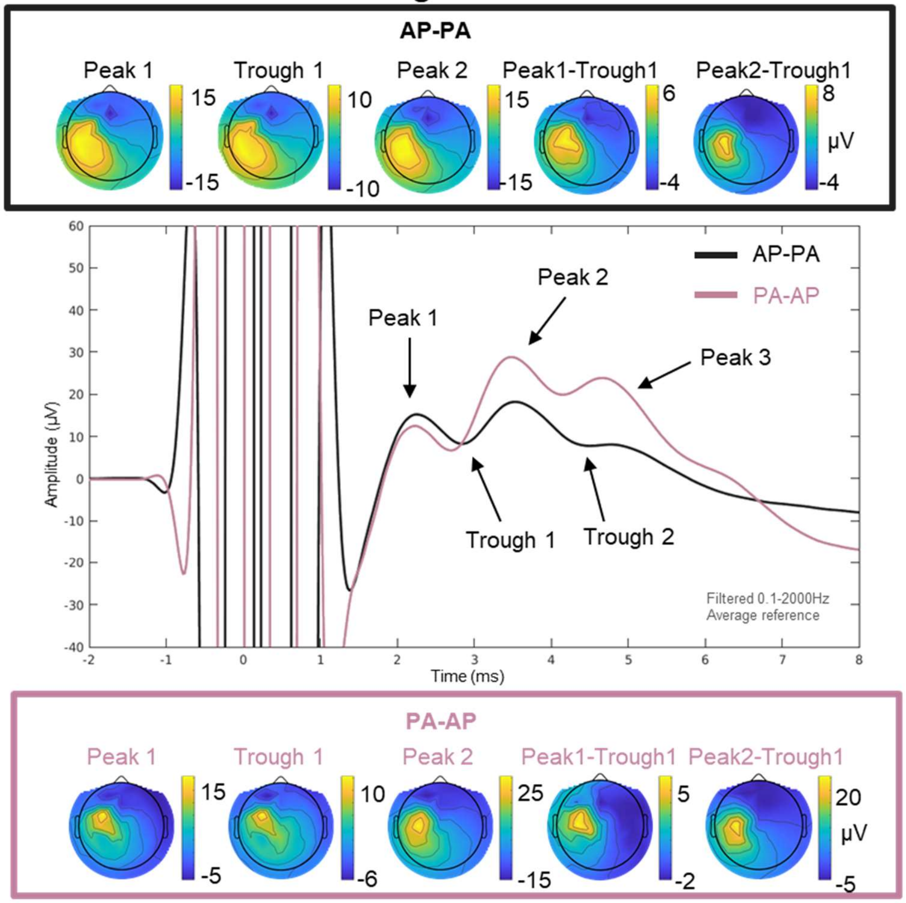

**Figure S10. Effects of re-referencing on i-TEPs.** (A) displays grand-averaged EEG butterfly plots for M1 TMS stimulation with AP-PA stimulation at 110%$RMT_{AP\text{-}PA}$ (black) and PA-AP at 110%$RMT_{PA\text{-}AP}$ (pink) stimulation using both the recorded forehead reference (top row) and re-referenced to average reference (bottom row). (B) Shows grand-averaged EEG response for an electrode close to the stimulation site and associated topographies for the same stimulation conditions as in (A). The i-TEPs still display similar properties regardless of the reference. For comparisons see Figure 1A in main text.
